# Do lanthanide-dependent microbial metabolisms drive the release of REEs from weathered granites?

**DOI:** 10.1101/2022.03.08.483559

**Authors:** Marcos Y. Voutsinos, Jacob A. West-Roberts, Rohan Sachdeva, John W. Moreau, Jillian F. Banfield

## Abstract

Prior to soil formation, phosphate liberated by rock weathering is often sequestered into highly insoluble lanthanide phosphate minerals. Dissolution of these minerals is critical for the release of phosphate to the biosphere, yet the microorganisms involved, and the genes required for lanthanide metabolism, are poorly understood. Here, we sampled weathered granite and associated soil to identify the zones of lanthanide phosphate mineral solubilization and genomically define the organisms implicated in lanthanide utilisation. We reconstructed 136 genomes from 11 bacterial phyla and found gene clusters implicated in lanthanide-based metabolism of methanol (primarily XoxF3 and XoxF5) are surprisingly common in microbial communities in moderately weathered granite where lanthanide phosphate minerals are dissolving. Notably, XoxF3 systems were found in Verrucomicrobia for the first time, and in Acidobacteria, Gemmatimonadetes, and Alphaproteobacteria. The XoxF-containing gene clusters are shared by diverse Acidobacteria and Gemmatimonadetes, and include conserved hypothetical proteins and transporters not associated with the few well studied XoxF systems. Given that siderophore-like molecules that strongly bind lanthanides may be required to solubilize lanthanide phosphates, it is notable that candidate siderophore biosynthesis systems were most prevalent in bacteria in moderately weathered rock, especially in Acidobacteria with lanthanide-based systems. We conclude that the confluence in the zone of moderate weathering of phosphate mineral dissolution, lanthanide utilisation, and methanol oxidation (thus carbonic acid production) may be important during the conversion of granitic rock to soil.

## Introduction

During the weathering of granite, a major component of Earth’s continental crust, some elements are redistributed from dissolved minerals into new, more stable ones, a critical process in the formation of soil (Kronberg and Nesbitt 1981). Microorganisms are key drivers of mineral weathering, but the mechanisms by which they promote mineral alteration, and the biogeochemical connections between microbial metabolisms and element redistribution are poorly understood.

A critically important component of granite weathering is the conversion of mineral-associated phosphate to bioavailable phosphorus. Apatite [Ca_5_(PO_4_)_3_(F,CI,OH)], the primary phosphate mineral, dissolves early, but much of the phosphorus released is sequestered into secondary phosphate minerals (Banfield and Eggleton 1989). The major cations in these secondary minerals are often lanthanides that may be released by dissolution of minerals such as allanite [(Ca,Ce,La)_3_(Fe^2+^,Fe^3+^)AI_2_O (Si_2_O_4_)(Si_2_O_7_)(OH)] and monazite [(Ca,Ce,La)POJ. The resulting minerals, such as rhabdophane (Ce,La,Nd,PO_4_·H_2_O) and florencite (La,Ce,Nd, Sm,Ba,Ca,Fe,Pb)AI_3_(PO_4_)_2_(OH)_6_, are exceedingly insoluble (<10^-25^*K*_sp_) (Firsching and Brune 1991; Gausse et al. 2016). This imposes a nutrient limiting effect on the ecosystem, commonly resulting in infertile soils.

Lanthanides were long considered to be of no biological relevance. However, now it is known that most facultative and obligate methylotrophic bacteria oxidise methanol to generate energy using XoxF enzymes (Keltjens et al. 2014), a class of lanthanide-dependent methanol dehydrogenases (MDH). These pyrroloquinoline quinone (PQQ) bound enzymes require lanthanides in their active site to catalyse the reaction of methanol to formaldehyde in the periplasm (Good et al. 2016). Genomic and metagenomic sequencing has revealed the large phylogenetic diversity of the XoxF family which is now believed to be the dominant MDH and evolutionary older than the more well studied calcium dependent MDH (MxaF) (Keltjens et al. 2014). To date, there are five known phylogenetically distinct clades of XoxF (XoxF1-5). XoxF5 and XoxF2 are the only experimentally studied clades, with representative sequences characterised as functional in the Alphaproteobacteria *Methylorubrum extorquens* AM1 (Skovran et al. 2011) and in *Verrucomicrobia fumariolicum* solV (Pol et al. 2014), respectively. Based on metagenomic studies of soil, XoxF3 is present in bacteria from a diverse range of phyla, including Proteobacteria, Acidobacteria, Gemmatimonadetes and Rokubacteria (Butterfield et al. 2016; Diamond et al. 2019). Sequence alignments demonstrated that these proteins have the conserved amino acid residues required for lanthanide-based functionality (Keltjens et al. 2014). XoxF5 is the largest clade reported to date, and together with XoxF1 and XoxF4, are only reported in Proteobacteria (Chistoserdova 2019).

Many experimental studies (Skovran et al. 2011; Pol et al. 2014; Deng, Ro, and Rosenzweig 2018; Ochsner et al. 2019) and one mini-review (Keltjens et al. 2014) indicate that most XoxF-based systems are comprised of the core periplasmic MDH XoxF, and homologs of XoxJ (Myung Choi et al. 2017) (a periplasmic binding protein of unknown function) and XoxG (Zheng et al. 2018) (a periplasmic membrane bound cytochrome specific to XoxF). XoxF3-based systems have not been experimentally studied, but one mini-review identified that a few XoxF3 operons include cytochrome genes (Cox, CtaG) (Keltjens et al. 2014). Overall, little is known about the suite of genes involved in XoxF-based systems outside of Proteobacteria.

Given that lanthanides in weathered granite are sequestered in highly insoluble minerals, theoretical considerations (Taunton, Welch, and Banfield 2000; Ochsner et al. 2019) and recent experimental work (Zytnick et al. 2022) suggest that specialised molecules such as siderophores may be required to induce lanthanide release. Siderophores are produced by biosynthetic gene clusters (BGCs) such as non-ribosomal peptide synthetases (NRPSs) and polyketide synthases (PKS) (Barry and Challis 2009). A recent preprint reports the first experimentally verified example of lanthanide-associated siderophore production in Alphaproteobacteria *Methyloruburm extorquens* AM1. This organism was shown to upregulate a Lanthanide Chelation Cluster (LCC) when supplied with poorly soluble Nd_2_O_3_ in vitro. LCC encodes a NRPS biosynthetic gene cluster containing a TonB dependent receptor and synthesises an aerobactin-like siderophore (Zytnick et al. 2022). This LCC is conserved across Methylobacterium species with some components present in other Alphaproteobacteria.

Interestingly, soils from which diverse lanthanide-based metabolisms have been inferred contain microbial communities that include organisms with numerous secondary metabolism gene clusters (Crits-Christoph et al. 2018)(Butterfield et al. 2016; Diamond et al. 2019; Sharrar et al. 2020)(Crits-Christoph et al. 2018). Thus, it is reasonable to speculate that the capacity to produce secondary metabolites that promote release of lanthanides from phosphate minerals may co-occur with genes for proteins that require lanthanides for functionality.

Despite many studies of microbial processes in soil (Waldrop, Balser, and Firestone 2000; Goldfarb et al. 2011; Pett-Ridge et al. 2013; Hultman et al. 2015; Malik et al. 2020), the microbial communities and their genomically-encoded functionalities in weathered rock have remained understudied. Nor has any study investigated the lanthanide-solubilizing capacity of microorganisms in such environments. The ‘onion skin’ feature of some granites, a concentric progression from weathered material towards freshly exposed rocks, provides an ideal opportunity to study both the mineralogy and microbiology along a weathering profile. Here, we characterised the mineralogical, geochemical and potential microbiological processes occurring in weathered I-type granite and associated soil, with a focus on potential for lanthanide utilisation and secondary metabolism. We report that diverse bacteria can perform lanthanide-based metabolism in moderately weathered rock, where lanthanide phosphates are solubilized.

## Results

### Sampling across a granite weathering profile

We sampled fresh and weathered I-type Burrumbeep granodiorite and soil from near Rocky Point Bushland Reserve (RPR), Ararat, Victoria, Australia **(Fig. 1).** The Victorian Geological Survey notes that this Ararat Suite granodiorite contains hornblende, biotite, zircon, apatite, allanite, sphene, calcite, fluorite, chlorite, quartz, and plagioclase and K-feldspars (Cayley and Taylor 2001). The mineralogy of the samples was confirmed via thin section analysis and scanning electron microscopy-based energy dispersive x-ray analysis (SEM-EDX). The densities of weathered and fresh rocks were measured and compared to provide an indication of the degree of alteration (i.e. mass loss). Samples with densities >2.5 g/cm^3^ were classified as nearly fresh rock, 2.4 to 2.2 g/cm^3^ as lightly weathered, 2.1 to 1.9 g/cm^3^ as moderately weathered, 1.8 to 1.6 g/cm^3^ as highly weathered and <1.6 g/cm^3^ as very highly weathered. Highly weathered samples lacking granitic texture were classified as soil. Nine samples were collected for metagenomics analysis, representing moderately weathered rock (1.9 g/cm^3^), highly weathered rock (saprolite; 20 cm below the surface) and soil.

**Figure 1.**
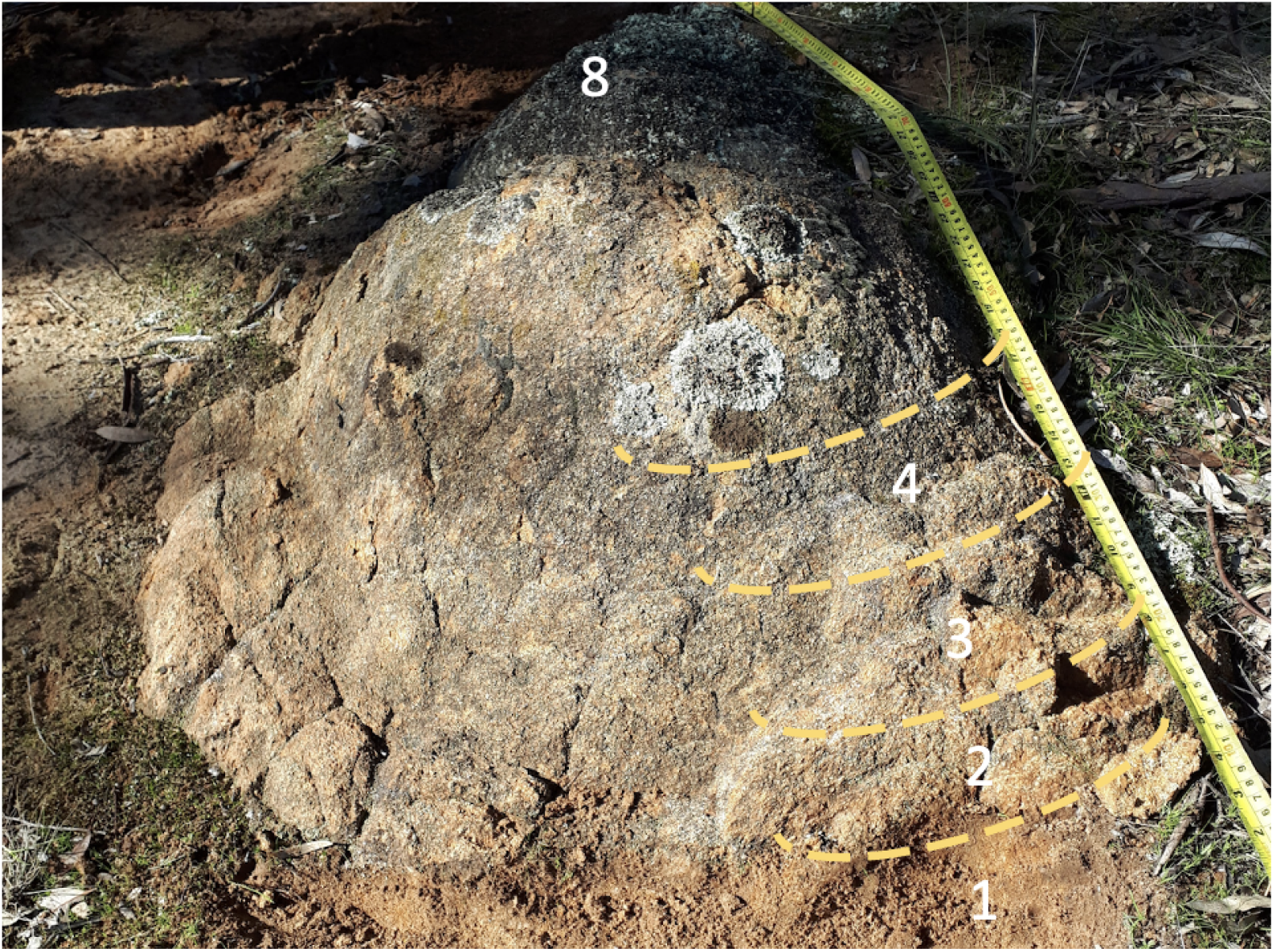
Scale photo of the exposed weathered I-type granite profile. Numbers indicate sampling locations for geochemical samples. Sample 1 is the most weathered sample and sample 8 is the least weathered sample. Photo taken after the first sample was collected.

### Microbial community composition as a function of weathering extent

We used ribosomal protein S3 (rpS3) gene sequences to assess the microbial diversity of the weathered profiles and used coverage of contigs carrying these genes to quantify relative microbial abundance. The rpS3 gene is a universal single-copy gene that is a good phylogenetic marker in soils because it assembles well from metagenomic data and is recovered more frequently than 16S rRNA genes (Olm et al. 2019). Across our metagenomic assemblies we identified 3,191 rpS3 sequences. All rpS3 sequences longer than 180 aa were grouped into 1,231 species groups (SGs) **(Supplementary Table 1)** based on 99% similarity (see methods). We classified all the rpS3 sequences by constructing a phylogenetic tree containing our sequences and the rpS3 sequences from diverse bacteria (Hug et al. 2016), and identified organisms from 18 phylum-level lineages **(Supplementary Data 1)** and their relative abundance **(Supplementary Table 2).** Actinobacteria and Alphaproteobacteria were the most abundant phyla in the moderately weathered region, while Actinobacteria and Acidobacteria were the most abundant in the soil and highly weathered regions. Some bacterial species groups exhibited high relative coverage while their phyla accounted for a small fraction of all the phyla represented in the community. This was most evident in the moderately weathered granite where sometimes only one microbial representative was present from Gemmatimonadetes, Acidobacteria, Chloroflexi and Verrucomicrobia despite exhibiting a high percentage of total coverage **(Fig. 2a).** This trend was also observed with Verrucomicrobia and Chloroflexi in the highly weathered region and Eremiobacterota, Chloroflexi, and Gemmatimonadetes in the soil. Ordination analysis **(Fig. 2b)** of the coverage data showed that communities sampled from the same weathering zone were more similar to each other than those from other weathered zones.

**Figure 2.**
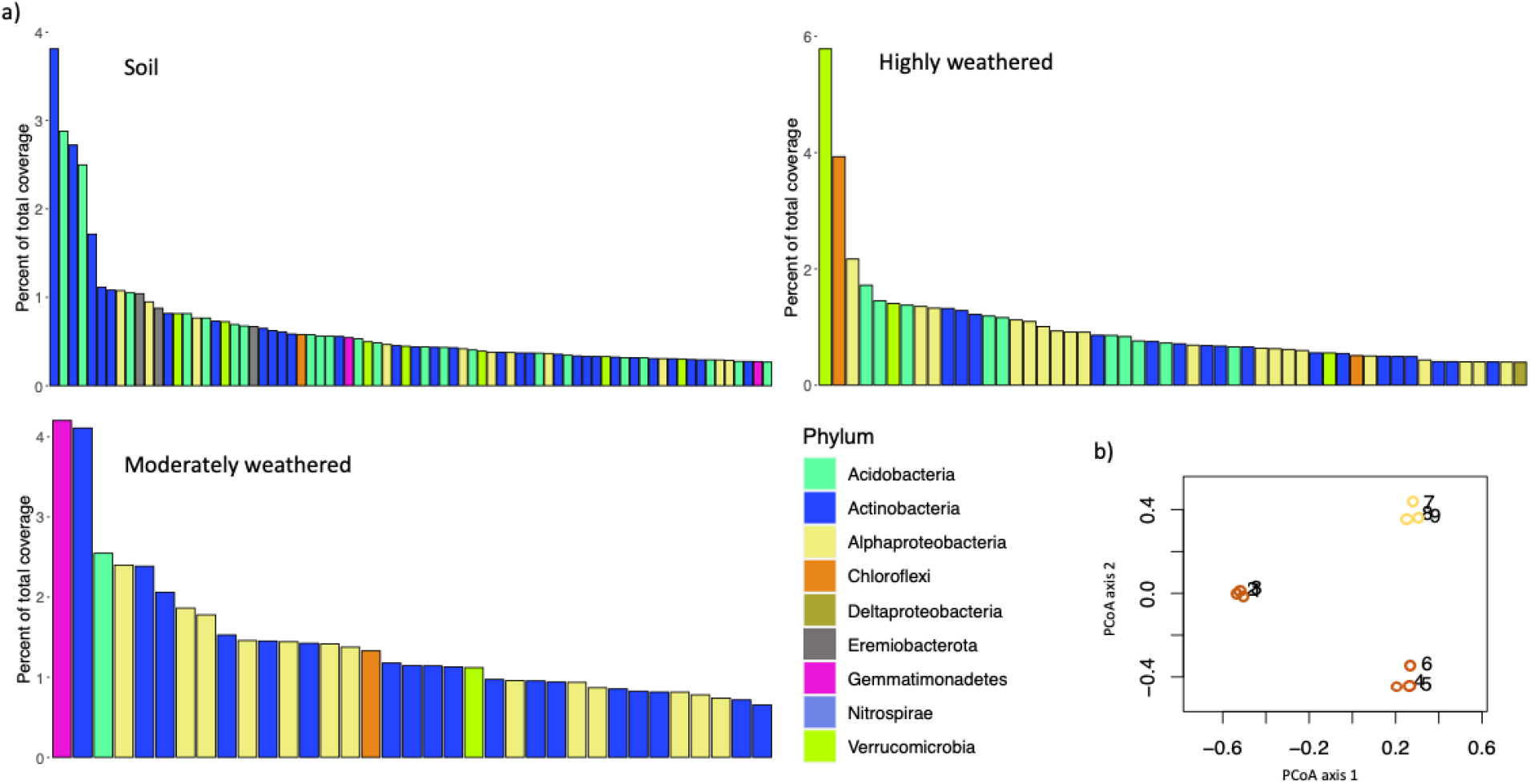
rpS3 species group (SG) diversity and abundance throughout a weathered granite profile, (a) percent of total relative coverage for the top 50% of the ribosomal protein S3 bearing contigs representing SGs (b) PCoA plot showing beta diversity of the nine communities collected from the soil (yellow), moderately (orange), and highly (red) weathered regions. Refer to Supplementary Table 1 for all SGs recovered and Supplementary Table 2 for their relative abundance.

### Abundance of the xoxF gene relative to degree of weathering

A customised HMM for PQQ-binding alcohol dehydrogenases was used to detect a dereplicated set of 411 methanol dehydrogenases encoded on the assembled contigs. All were of the XoxF type and contained the catalytic and cofactor binding residues required for activity, including those for PQQ, as well as conserved aspartate residues for binding lanthanides. No MxaF (Ca-dependent) representatives were identified. XoxF3 was most abundant, with 340 sequences, followed by XoxF5, with 63 sequences and 8 xoxF sequences that couldn’t be assigned to a clade. Overall, XoxF sequences were more commonly assembled in the moderately weathered (187) compared to highly weathered rock (54), and slightly more than in soil (170) **(Fig. 3 and Supplementary Table 3).** Thus, our findings extend our knowledge about XoxF in soils by showing that the capacity for lanthanide-dependent methanol oxidation is highly represented in diverse bacteria in weathered rock prior to its conversion to soil.

**Figure 3.**
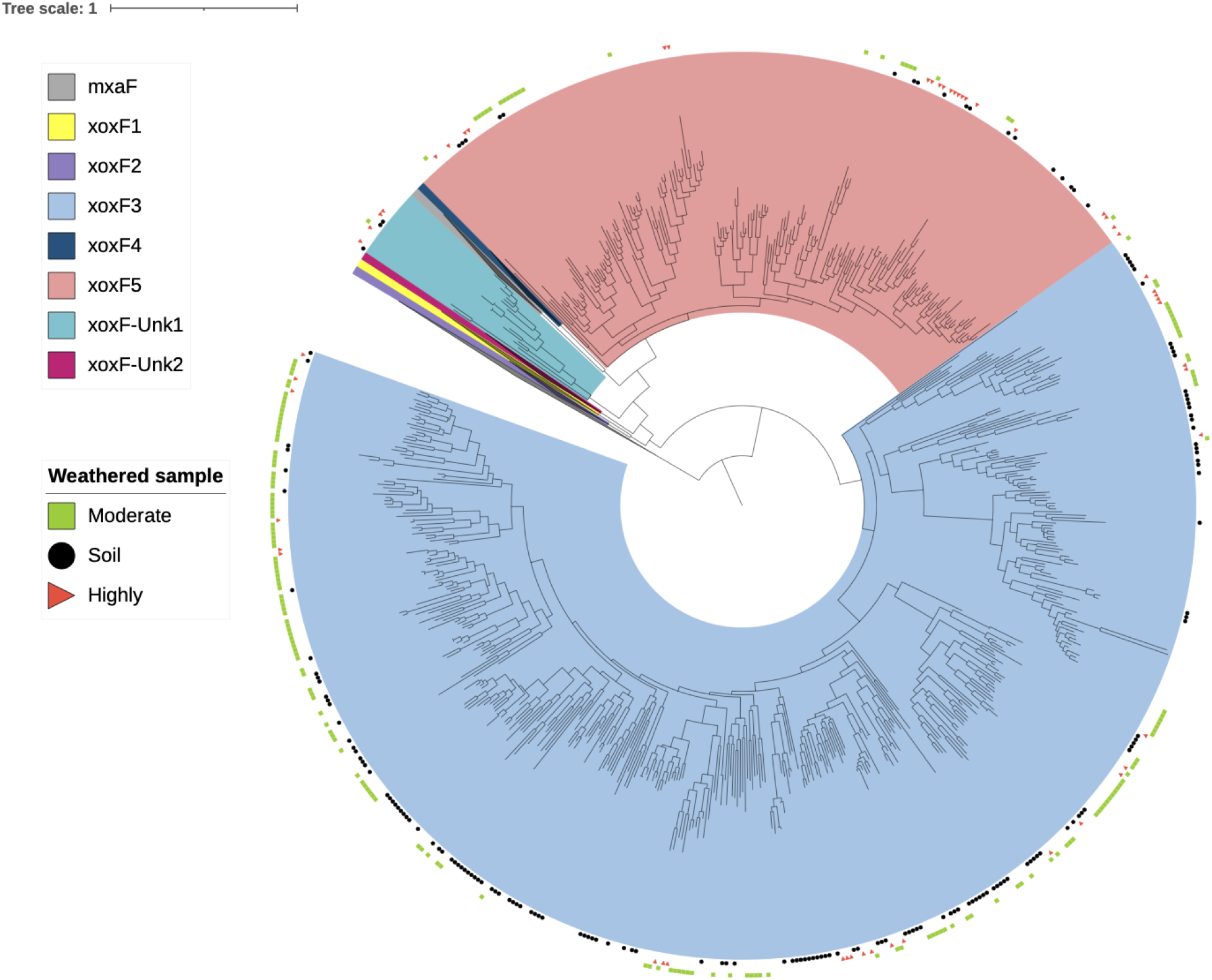
Phylogenetic analysis of the 411 methanol dehydrogenase sequences isolated from the metagenomes of the weathered granite profile. Weathered regions from which sequences were assembled are indicated by a green square (moderate), black circle (soil) or red right triangle (highly). Clades that do not contain sequences from this work are collapsed. Branches are coloured to indicate xoxF clade. Unknown subtypes indicate the references did not have a known clade. PQQ-ADH1 is present as an outgroup.

### Genome reconstruction

We reconstructed 136 non-redundant draft genome sequences (> 70% complete with <10% contamination) from 11 different bacterial phyla. The most frequently genomically sampled bacterial phyla was Actinobacteria, followed by Acidobacteria and Alphaproteobacteria **(Supplementary Fig. 1).** For reasons of their abundance and prior data indicating high lanthanide metabolism and secondary metabolite production capacity (Crits-Christoph et al. 2018; Diamond et al. 2019), we analysed in more detail the phylogeny of the organisms represented by 28 Acidobacteria genomes **(Supplementary Fig. 2).** A phylogenetic tree, using 16 ribosomal protein sequences of Acidobacteria genomes from this study and 150 reference genomes from (Diamond et al. 2019), representing many Acidobacteria groups, was built to classify the genomes. The majority were placed within Group 1 Acidobacteriales, Group 3 Solibacteres, and Group 4 Blastocatellia.

### XoxF3 diversity and phylogentic associations

In the set of 136 dereplicated genomes, 43 XoxF3 and 7 XoxF5 sequences were identified. XoxF was found in all regions but was much more common in the genomes of bacteria sampled from moderately compared to highly weathered rock and was also abundant in the genomes of soil bacteria. The XoxF genes were detected in Acidobacteria (gp 1 Acidobacteriia, gp 4 Blastocatellia, gp 2, and gp 3 Solibacteres), Gemmatimonadetes, Verrucomicrobia and Alphaproteobacteria genomes **(Fig. 4).** Interestingly, no XoxF were detected in Actinobacteria. All XoxF5 sequences occurred in Alphaproteobacteria genomes. We conclude that XoxF is utilised by bacteria from diverse phyla, is the dominant methanol dehydrogenase, and is particularly abundant in moderately weathered rock.

**Figure 4.**
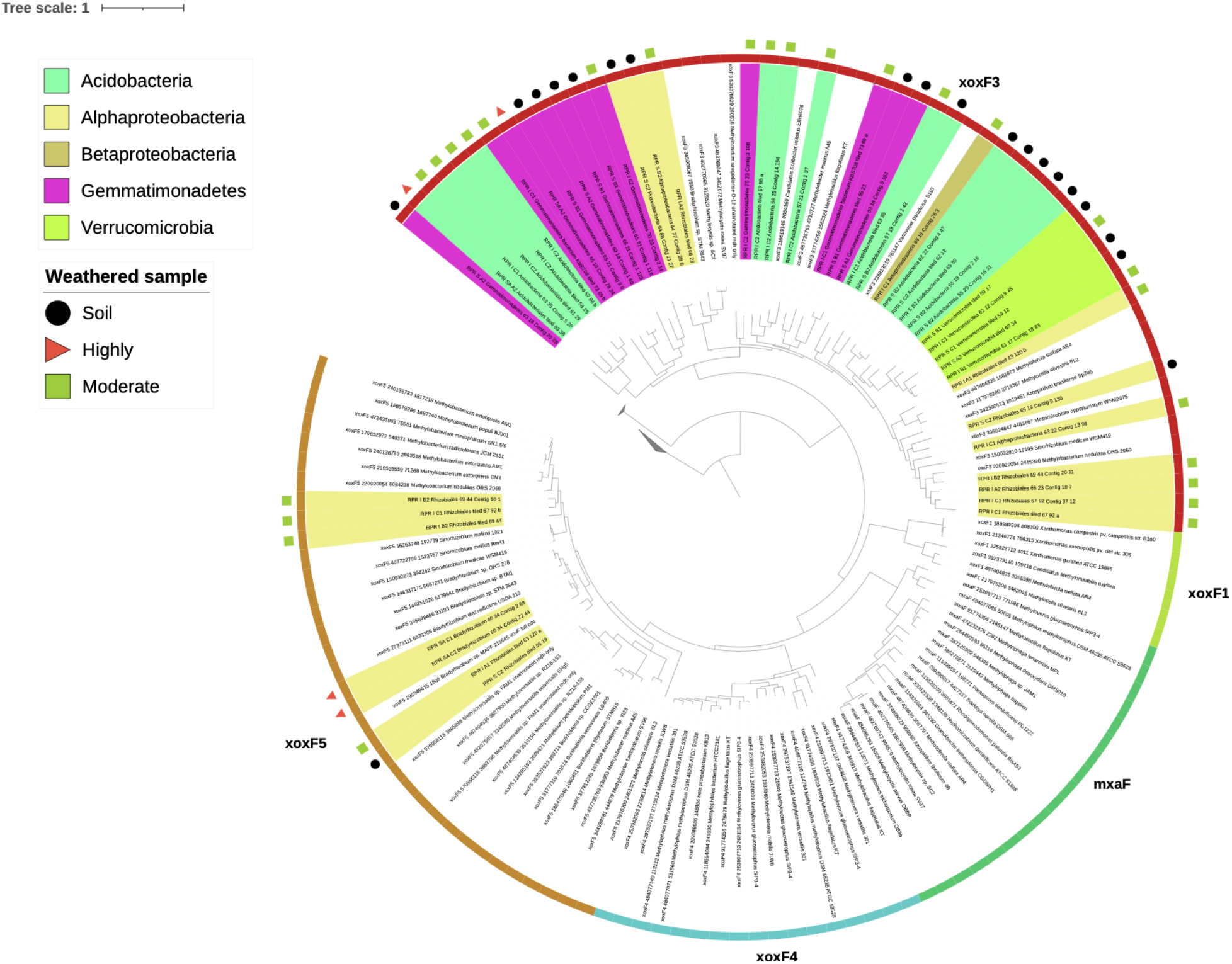
Phylogenetic analysis of the 50 xoxF sequences recovered from the dereplicated genome set and their associated taxonomy and weathered profile. Uncoloured names are reference XoxF sequences taken from (Diamond et al. 2019). Clades and tree rings are coloured to indicate xoxF type. Sample names are coloured to indicate their representative bacterial phylum. Tree was constructed using FastTree and visualised in iTOL.

### Finding of XoxF3 systems in Verrucomicrobia

Five high quality genomes from Verrucomicrobia contained XoxF3 systems **(Supplementary Fig. 3).** Upstream from XoxF3, all representatives contain a ~150 amino acid putative polyketide cyclase dehydrase enzyme. The enzyme contains a signal peptide and two transmembrane regions and has 50% percent identity to enzymes from primarily Methylobacterium. Two genomes also contain a cysteine-rich copper binding protein (DUF326). Other genes generally co-localising with XoxF3 include a mechanosensitive ion channel, a TonB dependent receptor, XoxJ (periplasmic binding protein of unknown function) and cytochrome enzymes including xoxG (XoxF specific cytochrome). To our knowledge, this is the first time a XoxF3 system has been predicted in bacteria of the phylum Verrucomicrobia.

### XoxF3 systems are conserved across Acidobacteria and Gemmatimonadetes

Five Gemmatimonadetes genomes contained 13 XoxF3 genes **(Supplementary Fig. 4)** with one genome (RPR_S_B1_Gemmatimonadetes_65_21) containing four XoxF3 genes. Almost all the XoxF3 systems contained xoxG, four contained xoxJ and two genomes contained a polyketide cyclase dehydrase. All Acidobacteria genomes with XoxF systems contained XoxF3 (12 genomes) co-localized with cytochrome genes (Coxl, Coxll, Coxlll, CtaG, CytC), xoxG, xoxJ and a small (−110 aa) hypothetical protein. Many genomes contained three small (~110-130 aa) hypothetical proteins surrounding a natural resistance-associated macrophage protein (NRAMP) domain **(Fig. 5).** Notably, seven Acidobacteria and two Gemmatimonadetes genomes contained NRAMP in the XoxF3 encoding region. Further, most Acidobacteria and two Verrucomicrobia and Gemmatimonadetes genomes contain XoxF3 systems including a −110-130 aa conserved hypothetical protein between CoxIII and CtaG. The Alphafold2 prediction of the 3D structure of this protein indicates high similarity between these proteins and vesicle-mediated transport proteins **(Supplementary Fig. 5).** To our knowledge, neither the NRAMP nor the putative vesicle-mediated transport proteins have previously been associated with XoxF systems.

**Figure 5.**
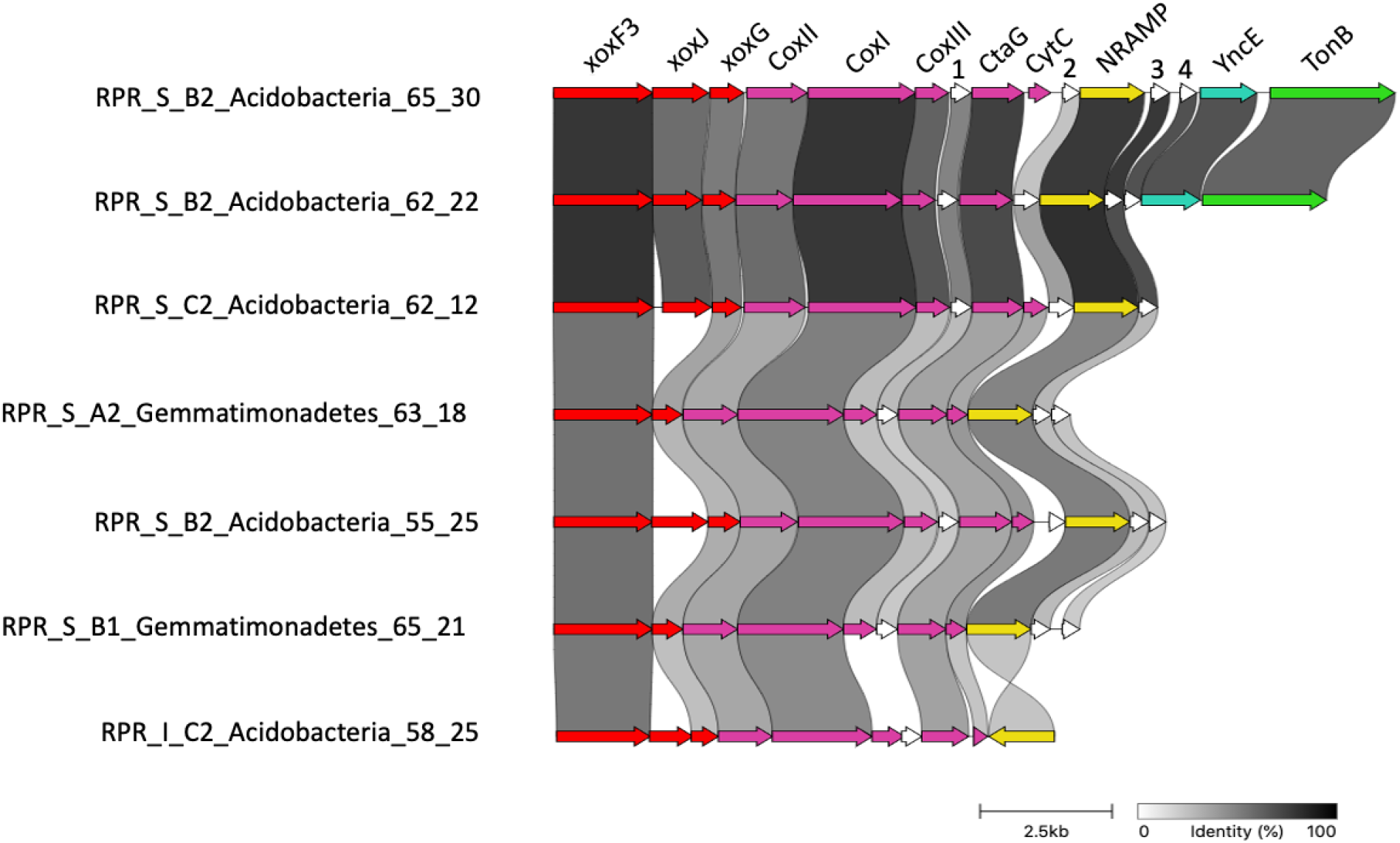
Gene cluster comparison of xoxF3 systems conserved across Acidobacteria and Gemmatimonadetes. Grey links show the percentage of identity between homologous proteins from different genomes. Abbreviations in order of appearance: xox, methanol dehydrogenase; Cox, cytochrome c oxidase; CtaG, cytochrome C oxidase assembly factor; CytC, cytochrome c; NRAMP, natural resistance macrophage protein; YncE, PQQ-dependent catabolism-associated beta-propeller protein; TonB, TonB dependent receptor.

### Lanmodulin and Pho regulon

Lanmodulin, a periplasmic lanthanide-binding protein that has been experimentally studied in the Alphaproteobacterium *Methylorubrum extorquens* (Cotruvo et al. 2018), was identified in one Bradyrhizobium (Alphaproteobacteria) genome but was not co-localized with XoxF. This protein contained 3 EF hand domains with the lanthanide specific proline residues (Cotruvo et al. 2018). We also identified another protein containing 3 EF hand domains but the domains contain neither the proline for lanthanide coordination nor the lysine residue for calcium coordination **(Supplementary Data 2).** We did not identify other homologs of lanmodulin in any other genomes or in the metagenomes in this study.

A complete Pho regulon was identified downstream from XoxF3 **(Supplementary Fig. 6)** in the genome of RPR_S_B1_Gemmatimonadetes_65_21. This regulon was identified in other Gemmatimonadetes genomes but genome fragmentation prevents clarification of the genomic context relative to the XoxF systems. Even this one case of close genomic association is interesting, given the close link between lanthanide release and phosphorus availability.

### Sources of lanthanides

An analysis of the extent of dissolution and pitting of apatite crystals enclosed within biotite showed that, as expected, increased degree of weathering correlated with increased extent of apatite dissolution **(Fig. 6a).** Secondary lanthanide phosphate minerals occurred as subhedral to euhedral crystals up to ~1 μm long and ~0.2 μm wide in hexagonal pits within biotite that were partly or previously occupied by apatite and as crystal aggregates on the surface of biotite **(Fig. 6b,c,d).** Secondary lanthanide phosphates were most abundant in the lightly weathered material, mostly dissolved in the moderately weathered material, and absent in the highly weathered material. This pattern was clarified by whole-rock elemental compositional data **(Supplementary Fig. 7 & Supplementary Table 4)** that showed that total lanthanide concentrations peaked in the lightly weathered (RPR1 - density 2.1 g/cm^3^) granite (427 ppm), where La concentrations were over five times higher than in fresh granite. The high enrichment of lanthanides in lightly weathered rock has been described previously (Banfield and Eggleton 1989; Voutsinos, Banfield, and Moreau 2021), and is attributed to redistribution of lanthanides from more weathered rock into phosphate phases at the weathering front.

**Figure 6.**
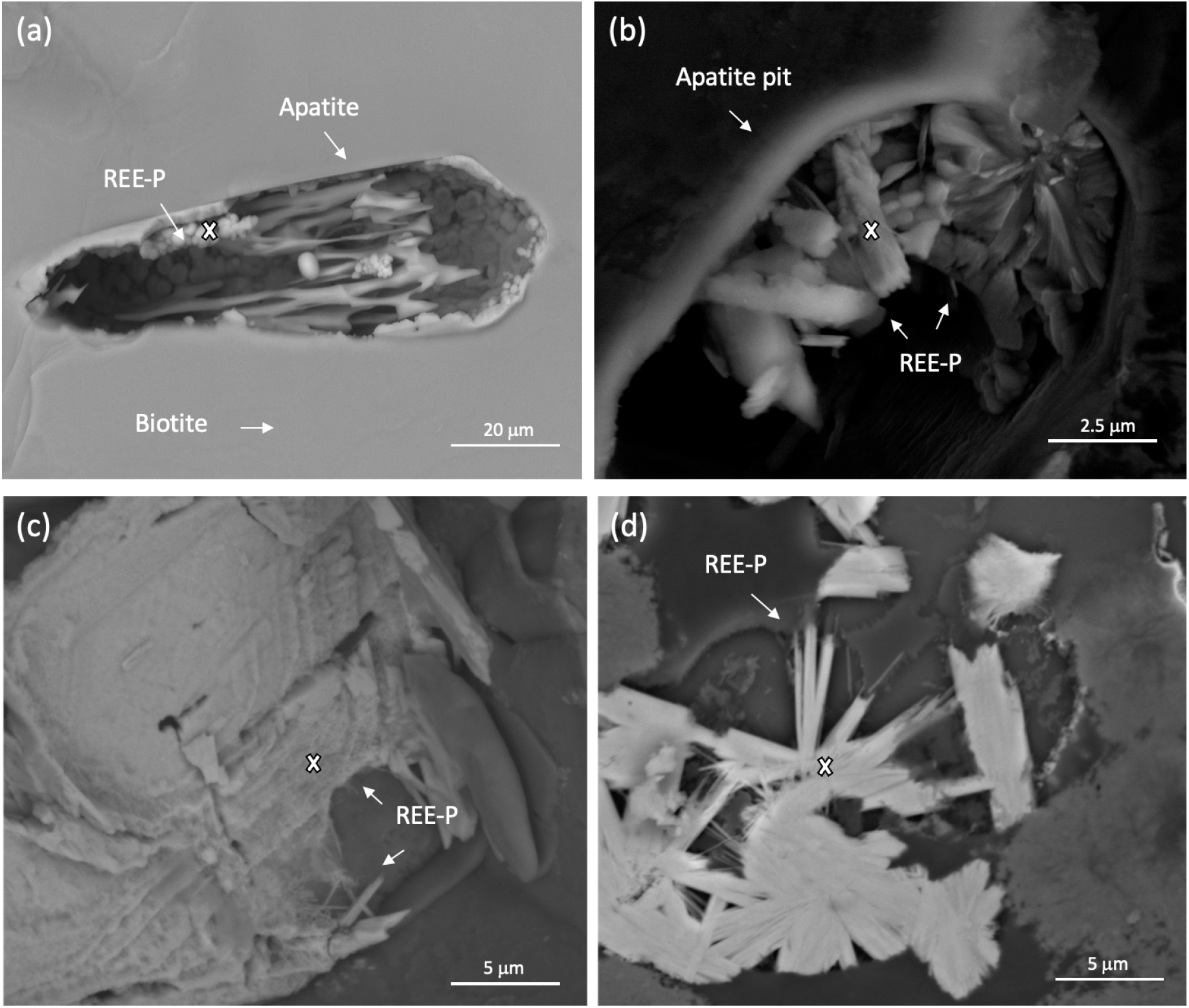
Scanning electron micrographs of secondary lanthanide phosphate minerals (a) replacing apatite (ei19 sp10), (b, c) precipitating in relict apatite pits (ei11 sp12 and ei29 sp47) and depositing on the surface of (d) biotite grains (ei33 sp 25). White ‘x’ indicates the site for EDX analysis. For EDX data, refer to Supplementary Table 6. SEM images captured at 10 kV.

Total lanthanide concentrations decrease substantially in the moderately weathered granite (264 ppm). Assuming approximately isovolumetric weathering (consistent with preservation of granitic texture), this supports the interpretation that lanthanides were lost much faster than other constituents in this zone. Some of the released La was likely transported into slightly weathered granite to account for the dramatic enrichment in this zone, yet lanthanides would be available for microbial utilisation. Concentrations were even lower in the highly weathered granite (214 ppm) and soil (80 ppm).

### NRPS/PKS abundance and distribution

Given that siderophore-like molecules are suspected to be important in promoting release of lanthanides, we investigated secondary metabolic capacity in the whole metagenomes (binned and unbinned sequences > 10 kbp). 1900 biosynthetic gene clusters were identified **(Supplementary Fig. 8 & Supplementary Table 5).** The amount of DNA sequence that encoded BGCs decreased from 0.53% in moderately weathered rock to 0.34% in the soil. This trend was paralleled by non-ribosomal peptide synthetases (NRPS, NRPS-like) and polyketide synthases (PKS types I, II, III and PKS-like). While the incidence of predicted siderophores (NRPS/PKS regions containing TonB dependent receptors) was highest in the soil compared to moderately and highly weathered rock. Two other transporters shown to be predictive of siderophore activity (Crits-Christoph et al. 2020), FecCD and Peripla_BP_2, did not occur in our samples.

We also predicted biosynthetic gene clusters in the set of 136 dereplicated genomes. We identified 457 biosynthetic gene clusters on contigs ≥10 kb and an additional 186 biosynthetic gene clusters on contigs less <10 kb **(Supplementary Table 6).** Of these, 168 were NRPS/PKS gene clusters, and they derived from genomes of bacteria from 10 different phyla (plus 54 smaller and possibly incomplete clusters). Biosynthetic gene clusters on contigs > 10 kb were most abundant in Acidobacteria and Actinobacteria, and most of these were NRPS/PKS systems **(Supplementary Fig. 9).** BCGs from genomes were most abundant in the moderately weathered granite while NRPS/PKS abundances were essentially the same in each of the sampled zones **(Supplementary Table 6).**

### Siderophore abundance and distribution

Transporters are required for the import and export of specialized biosynthesised metabolites such as siderophore-like molecules. We identified all classes of transporters across the 136 dereplicated genomes. The siderophore predictive TonB dependent receptor co-occurred with 10 NRPS/PVK biosynthetic gene clusters **(Table 1 & Fig. 7).** Three Acidobacteria (two group 1 Acidobacteriia and one group 2) genomes from the soil and moderately and highly weathered granite contained siderophores co-localized in genomes with XoxF3 systems. These XoxF systems occurred inside the antiSMASH predicted NRPS/PKS biosynthetic clusters. A further three Acidobacteria (two group 3 Solibacteres and one group 4 Blastocatellia) genomes from the moderately weathered granite contained XoxF and siderophore systems, but these were not co-localized with XoxF genes. Four genomes contain siderophore-like BGCs, but no XoxF systems were identified. Three of these genomes were Bacteroidetes, Sphingomonadales, Acidobacteria (group 1 Acidobacteriia) from the moderately weathered region and the other was for a Acidobacteria (group 4 Blastocatellia) from the soil. We did not observe any systems similar to the lanthanophore cluster of *Methylorubum Extorquens* AM1 (Zytnick et al. 2022).

**Figure 7.**
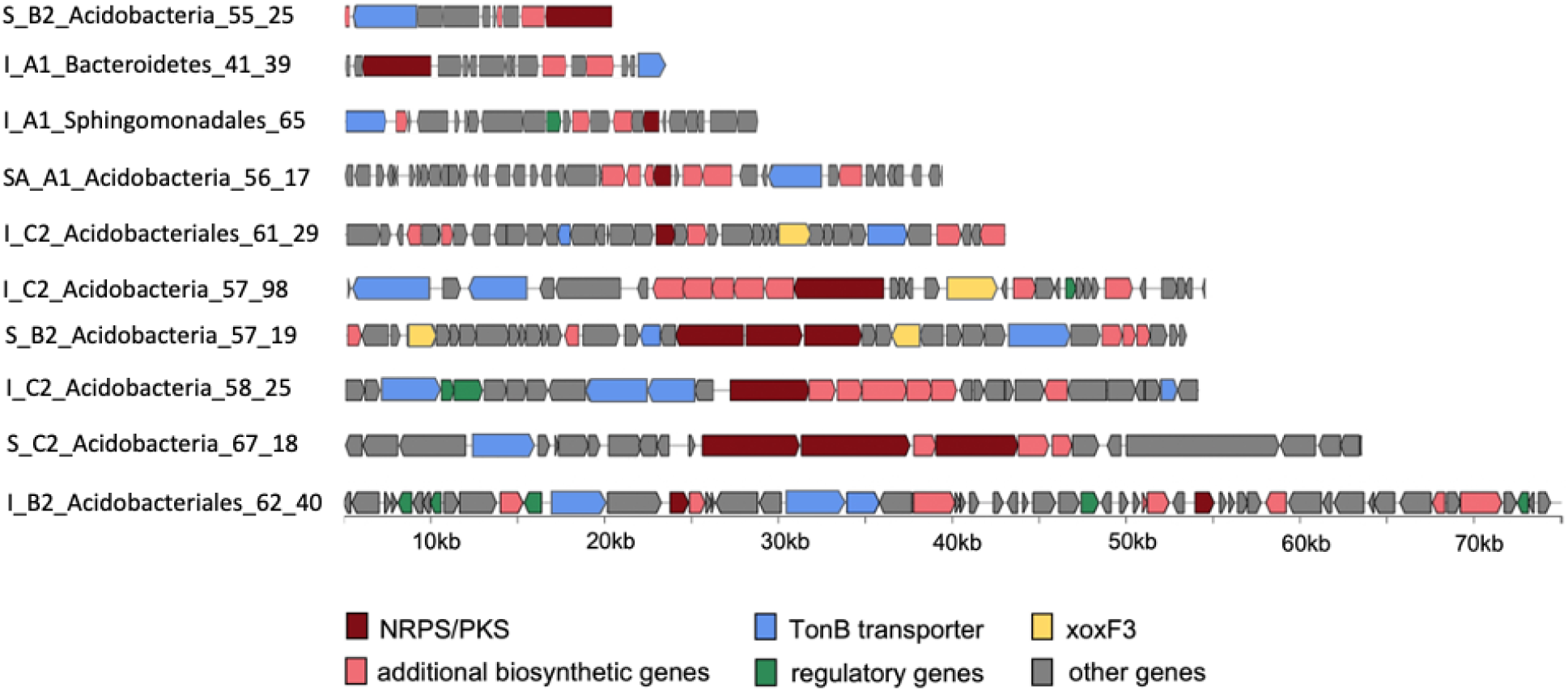
All biosynthetic gene clusters (BGCs) with predicted siderophore activity identified in the dereplicated genome set. All BGCs were detected using AntiSMASH 6.0. TonB transporters were identified using pfam_transporter20.hmm and xoxF sequences were detected using a customised HMM for PQQ-binding alcohol dehydrogenases (Diamond et al. 2019).

**Table 1.** Summary of 10 representative MAGs carrying xoxF3 and/or siderophores

| Weathered Region | MAG | Size (Mbp) | #Contigs | %Compl. | %Cont. | 16S | xoxF3 | Siderophore | Taxonomy |
| --- | --- | --- | --- | --- | --- | --- | --- | --- | --- |
| Moderate | Bacteroidetes_41_39 | 6.2 | 135 | 99.01 | 6.18 | - | - | + | Bacteroidia |
| Moderate | Sphingomonadales_65_36 | 2.34 | 54 | 92.45 | 3.11 | - | - | + | Alphaproteobacteria |
| Moderate | Acidobacteriales_62_40 | 3.74 | 16 | 78.42 | 0 | + | - | + | Gp 1 Acidobacteriia |
| Soil | Acidobacteria_67_18 | 6.08 | 221 | 89.16 | 5.13 | - | - | + | Gp 4 Blastocatellia |
| Moderate | Acidobacteria_57_98 | 4.84 | 36 | 97.39 | 0.43 | + | + | + | Gp 3 Solibacteres |
| Moderate | Acidobacteria_58_25 | 6.14 | 34 | 96.52 | 0.87 | - | + | + | Gp 3 Solibacteres |
| Soil | Acidobacteria_55_25 | 4.89 | 88 | 88.89 | 5.82 | - | + | + | Gp 4 Blastocatellia |
| Co-localization of xoxF3 and siderophore |  |  |  |  |  |  |  |  |  |
| Moderate | Acidobacteriales_61_29 | 3.24 | 22 | 94.68 | 3.85 | - | + | + | Gp 1 Acidobacteriia |
| Soil | Acidobacteria_57_19 | 5.16 | 163 | 80.34 | 8.55 | - | + | + | Gp 1 Acidobacteriia |
| High | Acidobacteria_56_17 | 3.44 | 31 | 77.99 | 5.37 | - | + | + | Gp 2 |

## Discussion

Soils have been a major focus of metagenomic studies that address microbial community composition (Diamond et al. 2019), community changes over depth and time (Butterfield et al. 2016; Sharrar et al. 2020), biosynthetic potential (Sharrar et al. 2020), microbial carbon compound processing (Woodcroft et al. 2018), response to climate change perturbation (Cheng et al. 2017; Viitamäki et al. 2021), viral ecology (Emerson et al. 2018) and many other aspects of biogeochemistry. However, microbial communities in weathered rocks - the precursors to soil - remain almost unstudied. Some exceptions include analysis of one metagenome of weathered shale (Lavy et al. 2019) and two metagenomes in a weathered granodiorite (Napieralski and Roden 2020). In the current study, we analysed the metagenomes collected through a granite weathering profile. This approach enabled us to predict microbial utilisation of lanthanides in organic compound oxidation and to provide clues regarding links among microbial metabolism, phosphate mineral dissolution and lanthanide and phosphate bioavailability.

Microorganisms preferentially colonise mineral surfaces (Hazen et al. 1991; Ohmura, Kitamura, and Saiki 1993; Ohmura et al. 1996) and can assist in the breakdown of silicate (Vandevivere et al. 1994; Ullman et al. 1996; Barker et al. 1998; Liermann, Barnes, et al. 2000; Santelli et al. 2001) and non-silicate minerals (Arnold, DiChristina, and Hoffmann 1988; Grantham, Dove, and Dichristina 1997; Hersman, Lloyd, and Sposito 1995; Welch, Taunton, and Banfield 2002; Davis, Nealson, and Lüttge 2007), thereby enhancing rock weathering. Iron is an abundant redox active component of some minerals and microbial iron oxidation can promote mineral breakdown (Napieralski and Roden 2020). Microorganisms also assist weathering via production of acids (Ullman et al. 1996). However, lanthanides can precipitate as insoluble secondary lanthanide phosphates such as rhabdophane, as reported previously (Banfield and Eggleton 1989; Taunton, Welch, and Banfield 2000; Sanematsu, Kon, and Imai 2015; Voutsinos, Banfield, and Moreau 2021) and as shown here. They are not solubilized via soil-associated acids (Gausse et al. 2016). Despite this, we find that the lanthanide-bearing XoxF-type methanol dehydrogenases are highly abundant in weathered rock and soil. In fact, the XoxF-based system was the only methanol dehydrogenase type identified at our site, indicating that any methanol oxidation occurring was via lanthanide-based, instead of calcium-based, metabolism. The high abundance of these enzymes in the weathering profile, along with microscopic evidence indicating loss of lanthanide phosphate phases, suggests that the lanthanides in enzymes derive from co-existing lanthanide phosphate minerals. This raises questions about the mechanisms responsible for lanthanide acquisition and uptake from the environment.

Microbes can access elements such as Fe, which is highly insoluble under oxic conditions, using siderophores, which have high affinity for ferric iron and are capable of solubilising iron oxide minerals (Hersman, Lloyd, and Sposito 1995; Liermann, Kalinowski, et al. 2000; Buss, Lüttge, and Brantley 2007). Long ago, the observation that soils were depleted in lanthanide phosphate minerals relative to underlying weathered rock motivated the suggestion that siderophores also may be required to dissolve lanthanide phosphate minerals (Taunton, Welch, and Banfield 2000). Here we show that lanthanide phosphates are solubilized in moderately weathered rock (as well as soil), where genes for lanthanide-based enzymes are abundant. A recent preprint describes a lanthanophore shown to induce poorly soluble Nd_2_O_3_ dissolution, and reports the first gene cluster implicated in this process (Good et al. 2016). The cluster is specific to *Methylorubrum extorquens* AM1 (Proteobacteria), and we find no related systems in bacteria with lanthanide-based enzymes in the weathered rock and soil studied here. Thus, we anticipate a variety of yet unknown siderophore-like molecules may perform the analogous function of complexing lanthanides to promote lanthanide phosphate mineral dissolution.

Siderophores are produced by large operonic sets of microbial biosynthetic gene clusters (BGCs). Genomes can contain numerous different BGCs, the products of which can be highly diverse, including siderophores, ionophores, antibiotics, antifungal compounds, and signalling compounds (Osbourn 2010). Typically, it is challenging or impossible to discern the types of molecules that BGCs produce by bioinformatics analysis. However, recent analyses of gene clusters with known products showed that siderophore products of BGCs can be predicted by identifying NRPS/PKS gene clusters that contain TonB dependent receptors (Crits-Christoph et al. 2020). We resolved Acidobacteria genomes containing XoxF3 systems within siderophore clusters, which suggests these siderophores may be involved in the solubilisation and transport of lanthanides into cells. We also identified Acidobacteria genomes containing both XoxF3 and siderophores systems, although they were encoded in different genomic regions. Despite this, these siderophores may also assist in lanthanide solubilisation and uptake, given that the lanthanide chelation cluster (LCC) of *Methylorubrum extorquens AM1* is located distantly from the XoxF machinery (Zytnick et al. 2022). The observation of genomes containing siderophore-like BGCs without XoxF systems in our site raises the possibility that some bacteria promote lanthanide phosphate mineral dissolution to access phosphorus, a byproduct of which is the release of lanthanides. Another possibility is that lanthanide-requiring microbes may be “cheaters” (Kramer, Özkaya, and Kümmerli 2020) and able to modify siderophore-like compounds produced by other organisms, optimising them for lanthanide binding. We speculate that the cyclase dehydrase (involved in polyketide synthesis) and acetyltransferase (GNAT) in the XoxF-encoding region of two Verrucomicrobia genomes may have such a function. These findings motivate experimental work to test these hypotheses.

To date, two transporters have been experimentally verified as required for lanthanide transport: the outer membrane siderophore-associated TonB-dependent transporter (Ochsner et al. 2019) and the periplasmic ABC-type transporter (Wehrmann et al. 2019). We identified two transport proteins not previously associated with XoxF systems, NRAMP and the mechanosensitive ion channel (MscS). The conserved co-occurrence of NRAMP with XoxF3 systems across Acidobacteria and Gemmatimonadetes indicate these genes may play a role in the transport of lanthanides. NRAMP proteins are involved in the transport of divalent metal cations such as Ca^2+^, Mn^2+^, Cd^2+^, Mg^2+^, and Fe^2+^ (Nevo and Nelson 2006). The five XoxF3 systems identified in Verrucomicrobia also contained a transporter channel not previously associated with XoxF systems. The MscS is a membrane channel that responds to cellular swelling when exposed to hypoosmotic solutions, by opening and allowing the cell to return to its resting volume (Hua et al. 2010). These transporters have also evolved to include potential roles in Ca^2+^ regulation (Cox et al. 2015). It would not be surprising if these channels play a role in lanthanide transport, given that Ca^2+^ and the lanthanides have similar atomic radii and the homology of calmodulin with lanmodulin (Cotruvo et al. 2018) and MxaF with XoxF (Pol et al. 2014). Given that these transporters are specific for metal cations rather than large organic molecules, and as lanthanide uptake involves a high affinity chelator (Ochsner et al. 2019), we suggest that these transporters uptake free lanthanides into the Verrucomicrobia, Acidobacteria, and Gemmatimonadetes cells.

Topsoils can contain higher biosynthetic capacity than deeper soils (Sharrar et al. 2020). This has been attributed to higher microbial interaction and competition in topsoils compared to deeper soils (Eilers et al. 2012; Brewer et al. 2019). However, this prior study did not include an analysis of weathered rock. In our study, microorganisms in the moderately weathered granite have the highest biosynthetic potential and this potential decreases with increased weathering. Further, we noted a higher incidence of NRPS/PKS BGCs in the moderately weathered rock relative to the heavily weathered rock and the soil. The pattern of siderophore-like BGCs decreasing with weathering zones is not supported by counts of siderophores with TonB transporters. However, we note that TonB are exclusive to gram negative bacteria, and that other transporters may be involved (Crits-Christoph et al. 2020). We infer that siderophores may be required to access critical Fe, lanthanides and P in the weathered rock.

In the current study, we sampled a spheroidal weathered granite outcrop, the surface of which was colonised by lichens **(Fig. 1),** assemblages of organisms well known to promote early-stage rock weathering (Chen, Blume, and Beyer 2000). Some organic compounds secreted by members of lichen communities are likely methylated, and thus a source of methanol (Diamond et al. 2019). Other sources of methanol include demethylation of pectin during growth of plant cells in surrounding vegetation and degradation of pectin and lignin (Kolb 2009). Regardless of the source, it is clear that methanol oxidation is an important microbial function during weathering of the granite studied here. Ultimately, oxidation of methanol in this zone leads to release of CO_2_ at the weathering front, and thus the production of carbonic acid. Therefore, lanthanide-using microbes, as well as heterotrophs whose growth is enabled by phosphate release, likely promote the dissolution of silicate minerals. The results reported here reveal how early rock colonisation may be linked to mineral weathering. A cascade of processes that result from phosphate and lanthanide element release will increase rock porosity and permeability, precursor steps for soil formation.

## Conclusion

Based on genome-resolved metagenomics conducted in mineralogical and geochemical context, we propose aspects of the complex biogeochemical processes of lanthanide acquisition, trafficking, and utilisation in diverse bacteria across a weathered granite transect. Specifically, we report conserved gene clusters in bacteria from several phyla that would be suitable for experimental studies to describe new lanthanide-based systems. We find that lanthanide utilising microorganisms are prevalent within the region where insoluble lanthanide-phosphate minerals dissolve and identify potential siderophore-like gene clusters that may be involved. Experimental characterization of these systems may lead to routes for improving access to phosphorus in P-limited agricultural soils and for recovery of lanthanides from economical resources for technological applications.

## Methods

### Sample locations and sample collection

We collected 5 samples for geochemical analysis representing fresh and weathered rock and associated soil from an exposed I-type granite outcrop near Rocky Point Road (RPR) located near Ararat, Victoria, Australia. Samples were collected inwards from the outer, most weathered material, sampling along ‘zones’ where possible towards the freshest material. Samples ranged in texture from soil to highly (saprolite), moderately, and lightly weathered material. The highly weathered material still retained their granitic texture. Six months later we collected 9 samples for biological analysis around the same profile representing soil, highly weathered (saprolite) and moderately weathered granite. The geochemistry of these samples was also analysed. Samples were collected using a metal hand trowel sterilised using ethanol and flame. In the field, immediately after collecting the material, samples were homogenised, placed into sterile bags and flash frozen in a mixture of dry ice and ethanol and placed into an esky with dry ice for transport to the laboratory. Samples were delivered to the laboratory the same day and stored at −80C before DNA extraction.

### Mineralogical sample preparation and analysis

Given that apatite is the most likely source for P required for REE/ P-bearing mineral precipitation, we focused on the surfaces of apatite crystals and relict apatite pits to locate secondary lanthanide phosphate minerals. Apatite crystals were identified in biotite grains with their c axes oriented parallel to the basal plane of biotite. As biotite contained euhedral apatite crystals up to 100 μm long, biotite grains were extracted from the weathered granite samples using tweezers under a binocular microscope and split along their cleavage plane using a scalpel to reveal interior basal planes. Biotite grains were used for apatite and secondary lanthanide phosphate characterisation using SEM energy-dispersive X-ray spectroscopy (EDX). Cleaved biotite grains were mounted on SEM stubs and glass slides using double-sided carbon adhesive, carbon-coated and secondary lanthanide phosphates were characterised using a FEI Teneo VolumeScope. Given their high average atomic number, REE/P-bearing minerals were located using back-scattered electron imaging (BSE), at an accelerating voltage of 10 kV. At least 50 biotite grains were examined per sample with anywhere from 0-30 secondary REE/P minerals analysed per sample. Mineral chemistry was determined using EDX analysis; was semi-quantitative and standardless with a predicted error rate of at least 10%. Minerals phase analysis was repeated and reproduced when possible to reduce experimental error. Sample density was determined by weighing samples, then coating them with parafilm and weighing samples in water.

### Whole rock chemical analysis

Fist sized rock samples were crushed using a rock crusher and then further milled into a powder using an agate ring mill. Whole-rock elemental analysis was performed using the Applied Technologies 7700 ICP-MS instrument (see Supplementary Materials for detailed methods) with an expected error rate of −5%.

### DNA extraction, sequencing and metagenomic assembly

DNA was extracted from 10 g of each sample using the PowerMax Soil DNA isolation kit (MoBio Laboratories). Metagenomic library preparation and DNA sequencing were performed at the Next Generation Sequencing Facility, Western Sydney University. Metagenomic libraries were prepared using the IDT Lotus PCR-free kit and sequencing was performed on a NovaSeq 6000 platform, producing 250 bp paired-end reads. From 9 samples 18 metagenomes were produced and were processed individually. Raw reads were trimmed of adapters using bbduk (https://sourceforge.net/projects/bbmap/) with the following parameters: reference=Contaminants/adapters.fa k=23 mink=11 hdist=1 tbo tpe ktrim=r ftm=5. The reads were also screened for Phix and Illumina artifacts using bbduk and the following parameters: reference=resources/phix174_ill.ref.fa.gz,Contaminants/lllumina.articfacts.2013.12.fa.gz k=31 hdist=1. Finally, reads were quality trimmed using Sickle (https://github.com/najoshi/sickle) with the following parameters:: pe -q 20 -I 20. The samples were individually assembled using megahit with default parameters.

### Metagenome annotation

All samples were filtered to remove contigs smaller than 1000 bp using pullseq (https://github.com/bcthomas/pullseq). Open reading frames (ORFs) were predicted on all contigs using Prodigal v2.6.3 (Hyatt et al. 2010) with the following parameters: -m -p meta. Predicted ORFs were annotated using USEARCH (Edgar 2010) to search all ORFs against Uniprot (“UniProt: The Universal Protein Knowledgebase” 2017), Uniref90 and KEGG (Kanehisa and Goto 2000). 16S ribosomal rRNA genes were predicted using the 16SfromHMM.py script from the ctbBio python package using default parameters (https://github.com/christophertbrown/bioscripts). Transfer RNAs were predicted using tRNAscan-SE. The metagenomes and their annotations were then uploaded to ggkbase (https://ggkbase.berkeley.edu).

### Genome binning, filtering and dereplication

Metagenome assemblies were binned using differential coverage binners MaxBin2 (Wu, Simmons, and Singer 2016), MetaBAT (Kang et al. 2015) and VAMB (Nissen et al. 2021). The highest quality bins from each metagenome were selected via DasTool (Sieber et al. 2018). Bins quality was manually assessed using the ggkbase platform based on contig coverage, GC values, and the inventory of 51 bacterial single-copy genes. Bin completeness and contamination was also analysed using CheckM (Parks et al. 2015) lineage_wf using a threshold of >70% completeness and <10% contamination. Finally, bins were dereplicated at 98% nucleotide identity using dRep (Olm et al. 2017).

### Ribosomal protein S3 (rpS3), clustering and diversity analysis

All proteins predicted from the 18 metagenomes were analysed for rpS3 genes using a custom hidden Markov model (HMM) from (Diamond et al. 2019) with a threshold score of 40. Across all metagenomes we identified a total of 3,191 rpS3 sequences passed the assigned HMM threshold. Only rpS3 sequences with lengths of 180 to 450 amino acids were included resulting in 2,181 rpS3 proteins. We then clustered the sequences at 99% ID using USEARCH to obtain clusters that approximately equate to species-level identification which we refer to as species groups (SGs). The following USEARCH command was used: -cluster_fast RPR_rpS3_filtered_seqs.faa -sort length -id 0.99 -maxrejects 0 -maxaccepts 0 -centroids RPR_rpS3_filtered_seqs_centroids.faa. This resulted in 1,230 dereplicated rpS3 proteins, each approximately representing a species group. We then mapped the reads from each sample to the rpS3 bearing scaffold using BBMap for abundance quantifications of each SG. BBMap (http://sourceforge.net/projects/bbmap/) was then used to calculate the average coverage per base pair of each SG. The coverage was then normalised to percent abundances in each sample. rpS3 SGs were classified at the phylum level by constructing a phylogenetic tree containing our sequences and rpS3 references taken from ref (Hug et al. 2016). Our 1,230 rpS3 sequences were concatenated with the reference set and aligned using FAMSA. The resulting alignment was stripped of columns containing >90% gaps using trimal and a phylogenetic tree was constructed from the alignment using FastTree. Sequences were then manually classified to the phylum level based on their position relative to reference sequences in the tree. A principle coordinate analysis (PCoA) was performed using the Bray-Curtis distance measure which was calculated using the R programming tool and the vegan package.

### Genome phylogenetic classification

To taxonomically classify the microorganisms represented by the 136 dereplicated bins we used the combination of a concatenated ribosomal protein tree and a rpS3 protein tree. For the ribosomal protein tree, we searched each genome for 16 ribosomal proteins (RP16) using GOOSOS.py (https://github.com/jwestrob/GOOSOS). The following HMMs were used: Ribosomal_L2 (K02886), Ribosomal_L3 (K02906), Ribosomal_L4 (K02926), Ribosomal_L5 (K02931), Ribosomal_L6 (K02933), Ribosomal_L14 (K02874), Ribosomal_L15 (K02876), Ribosomal_L16 (K02878), Ribosomal_L18 (K02881), Ribosomal_L22 (K02890), Ribosomal_L24 (K02895), Ribosomal_S3 (K02982), Ribosomal_S8 (K02994), Ribosomal_S10 (K02946), Ribosomal_S17 (K02961), and Ribosomal_S19 (K02965). Ribosomal S10 model PF00338 was also used for identification of Chloroflexi. A total of 120 genomes containing at least 8 ribosomal proteins on a single contig were included. Our ribosomal protein sequences were then individually aligned using FAMSA and concatenated using the concatenate_and_align.py script from GOOSOS (https://github.com/jwestrob/GOOSOS/blob/master/Concatenate_And_Align.py). The resulting alignments were stripped of columns containing 90% gap positions using Trimal (Capella-Gutiérrez, Silla-Martínez, and Gabaldon 2009) with the parameter-gt 0.1. A phylogenetic tree was constructed using IQ-TREE and the following settings: iqtree -s RPR_RP16.mfaa -bb 1000 -nt AUTO -ntmax 48 -mset LG+FO+R. Genomes were then classified at the phylum level using GTDB-TK. If a genome was not included in the ribosomal protein tree, its taxonomy was determined via the rpS3 tree. For Acidobacteria genomes, class-level lineages were determined by building a ribosomal protein tree with 150 Acidobacteria reference genomes from (Diamond et al. 2019) and manually classified based on their position relative to reference sequences in the tree.

### XoxF identification and classification

For methanol dehydrogenase (XoxF) identification and classification of clades we constructed a phylogenetic tree to discriminate homologous, but functionally distinct proteins that can’t be identified by HMM search alone. XoxF sequences were identified in metagenomes using a custom HMM for PQQ-binding alcohol dehydrogenases taken from (Diamond et al. 2019). Across all metagenomes we identified 927 XoxF sequences. Proteins greater than 300 amino acids in length were retained and dereplicated at 95% similarity using CD-HIT resulting in 411 XoxF sequences. These sequences were concatenated with a reference set (Lavy et al. 2019; Diamond et al. 2019) and aligned using MAFFT using the following parameters: --localpair --maxiterate 1000 --reorder. The gaps were then removed from the alignments using trimal. A phylogenetic tree was constructed using FastTree and sequences were manually classified based on their relationship with the XoxF reference set. The above method was repeated for the XoxF sequences derived from the dereplicated genome set resulting in 49 XoxF sequences. Regions of conservation in XoxF systems between bacterial genomes were identified using the progressive mauve genome algorithm in Geneious with default settings and the clinker gene cluster comparison tool (Gilchrist and Chooi 2021).

### Biosynthetic gene cluster (BGC) and siderophore prediction

To identify biosynthetic gene clusters (BGCs), antiSMASH 5.0 (Blin et al. 2019) was run on the metagenomes and the final set of dereplicated genomes using default parameters. BGCs retrieved from the metagenomes were dereplicated using CD-HIT at 95% and normalised to percent abundance in each sample. Only BGCs on contigs greater than 10 kb were included in the analysis from both the genomes and the metagenomes. The antiSMASH tool only classifies BGCs as siderophores when they contain lucA/lucC genes which are specific for aerobactin and aerobactin-like siderophores. To predict the occurrence of siderophores outside of aerobactin we then ran two Pfams on the BGCs pfam_transporter20.hmm and all_sbp.hmm to identify the the BGCs that contain the transporters: FecCD, Peripla_BP_2, and TonB_dep_Rec. Previous work has shown these transporters are predictive of siderophore activity (Crits-Christoph et al. 2020).

## Supplementary Data

**Supplementary Figure 1.**
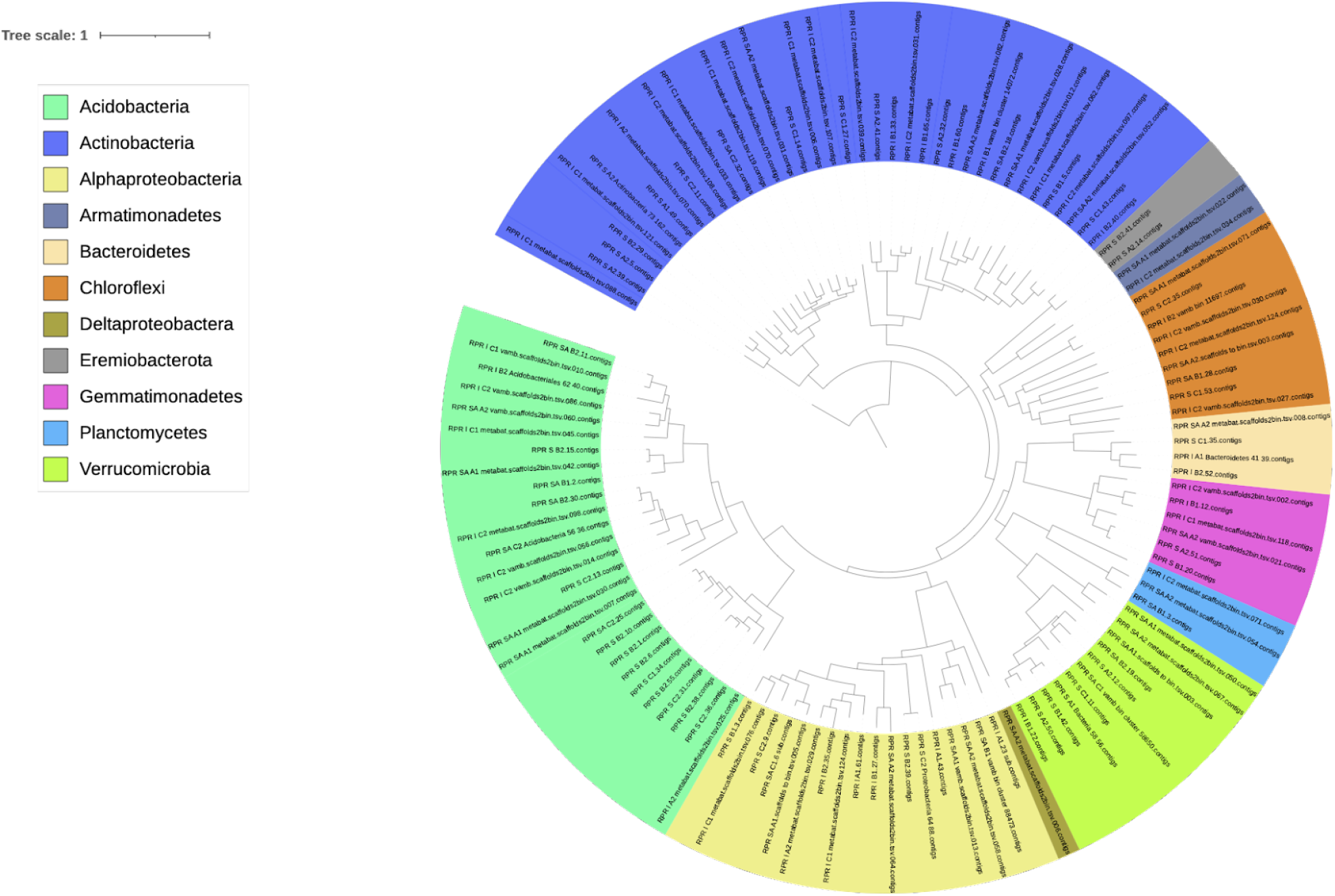
Phylogenetic tree constructed with a concatenated alignment of 16 ribosomal proteins from 120 bacterial genomes resolved from this study. Genome names are coloured according to their bacterial phylum.

**Supplementary Figure 2.**
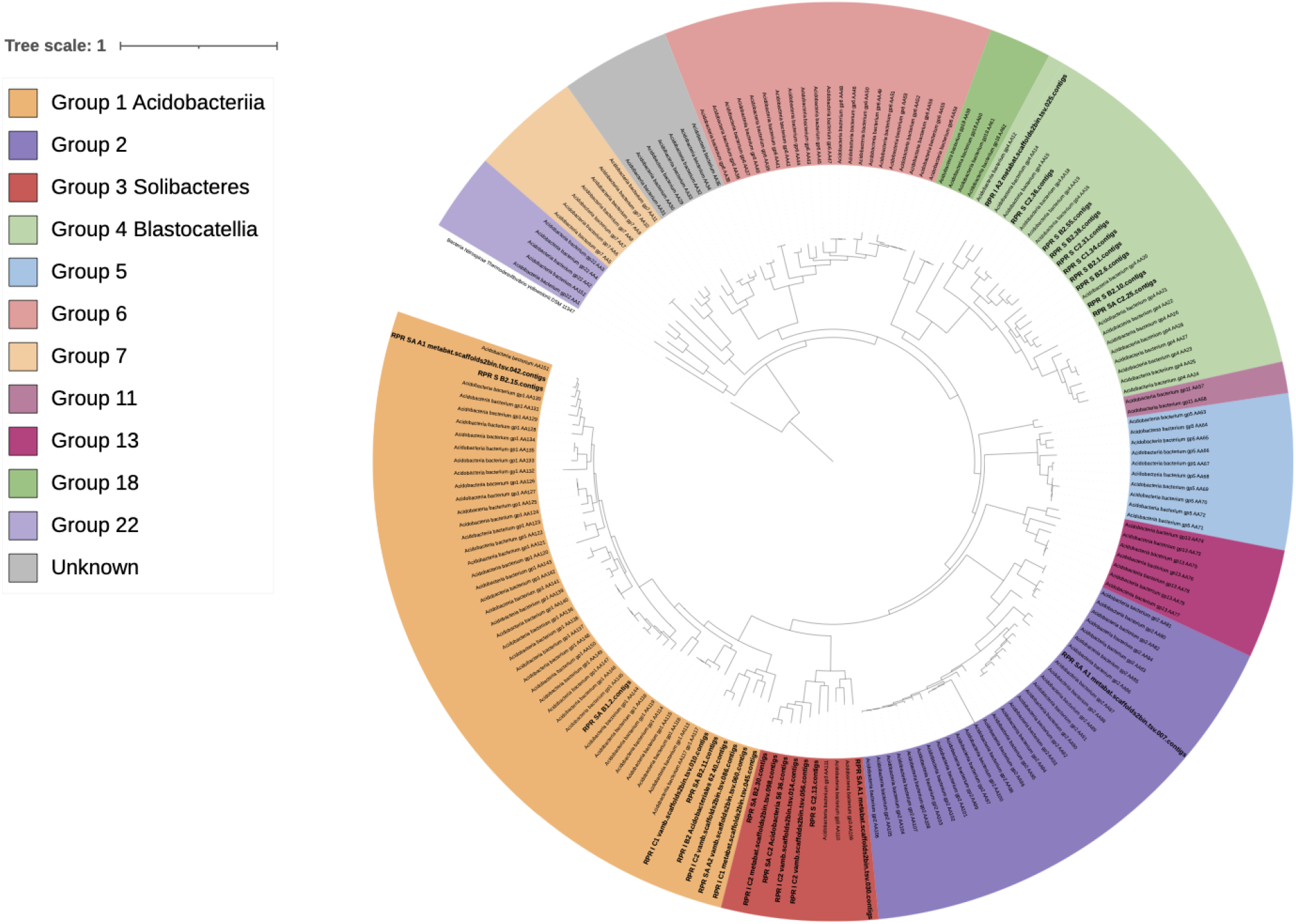
Phylogenetic tree constructed with a concatenated alignment of 16 ribosomal proteins. The tree includes 28 acidobacteria bacterial genomes where 8 or more ribosomal proteins were identified from this study (bold) and 150 acidobacteria reference genomes from (Diamond et al. 2019). Nitrospira is present as an outgroup.

**Supplementary Figure 3.**
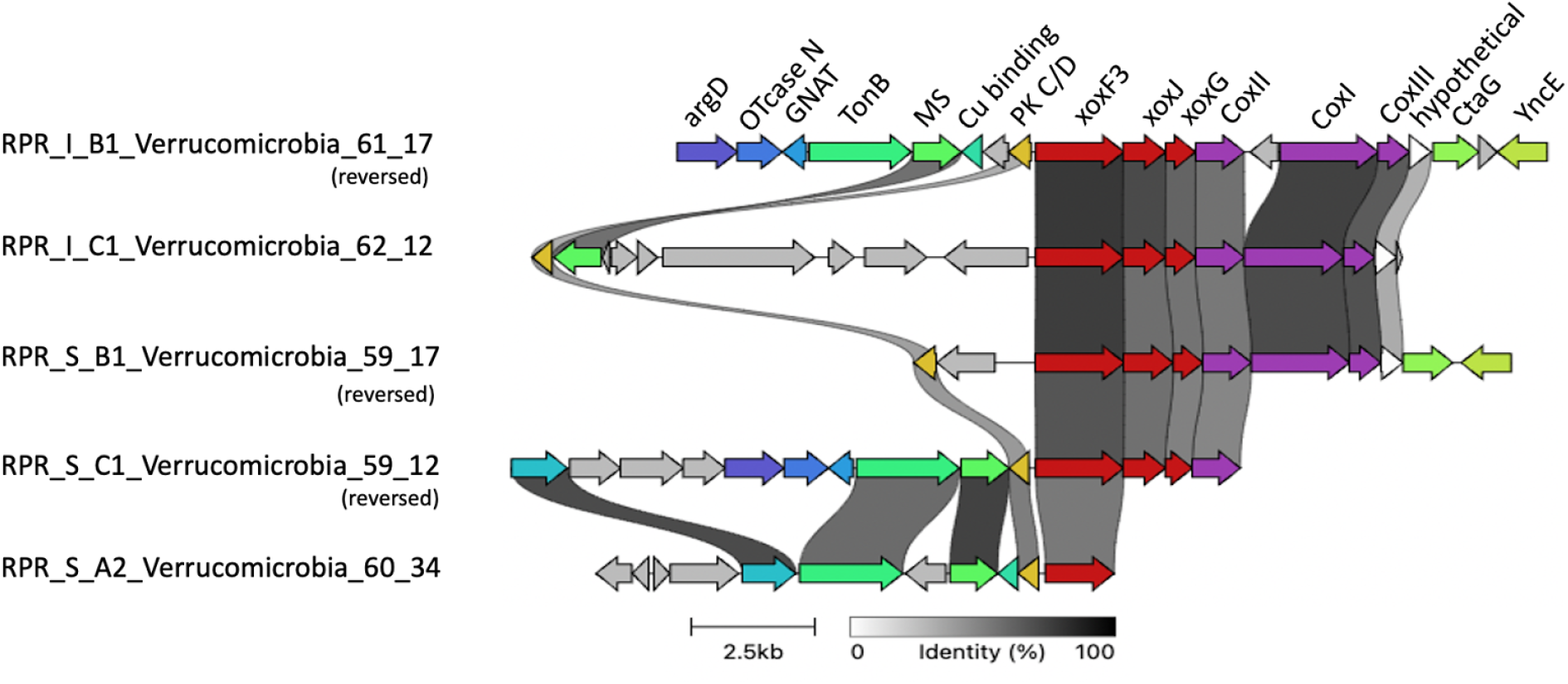
Gene cluster comparison of xoxF3 systems from Verrucomicrobia genomes. Grey links show the percentage of identity between homologous proteins. Genes appearing once are grey and appearing twice or more are colour coded. Abbreviations in order of appearance: argD, acetylornithine aminotransferase; OTnase N, ornithine carbamoyltransferase; GNAT, N-acetyltransferase; TonB, TonB dependent receptor; MS, mechanosensitive ion channel; Cu binding, copper binding; PK C/D, polyketide cyclase dehydrase; xox, methanol dehydrogenase; Cox, cytochrome c oxidase; CtaG, cytochrome C oxidase assembly factor; YncE, PQQ-dependent catabolism-associated beta-propeller protein.

**Supplementary Figure 4.**
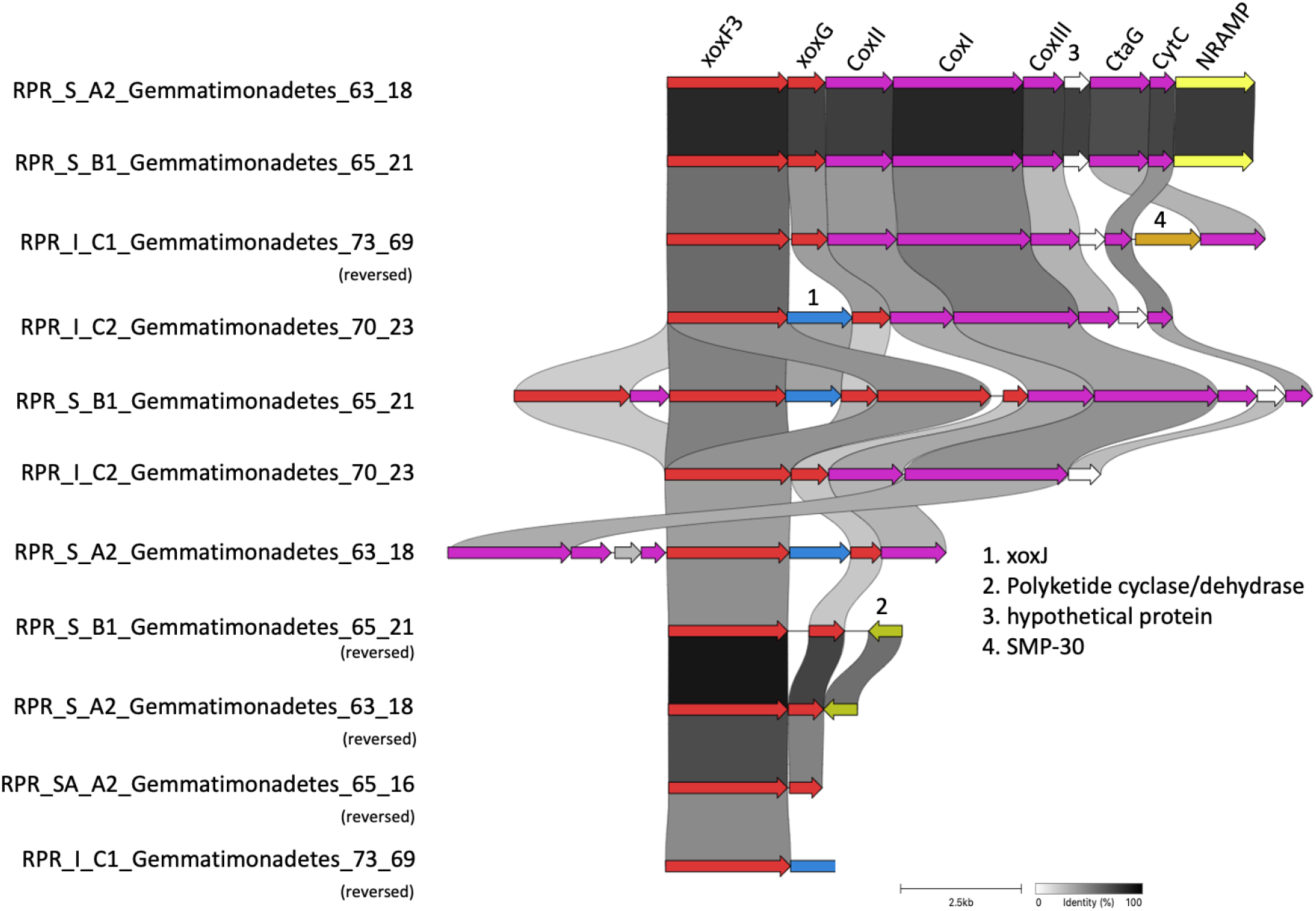
Gene cluster comparison of xoxF3 systems from five Gemmatimonadetes genomes. Grey links show the percentage of identity between homologous proteins. Abbreviation in order of appearance: xox, methanol dehydrogenase; Cox, cytochrome c oxidase; CtaG, cytochrome c oxidase assembly factor; CytC, cytochrome c; SMP-30, senescence marker protein 30; NRAMP, natural resistance macrophage protein.

**Supplementary Figure 5.**
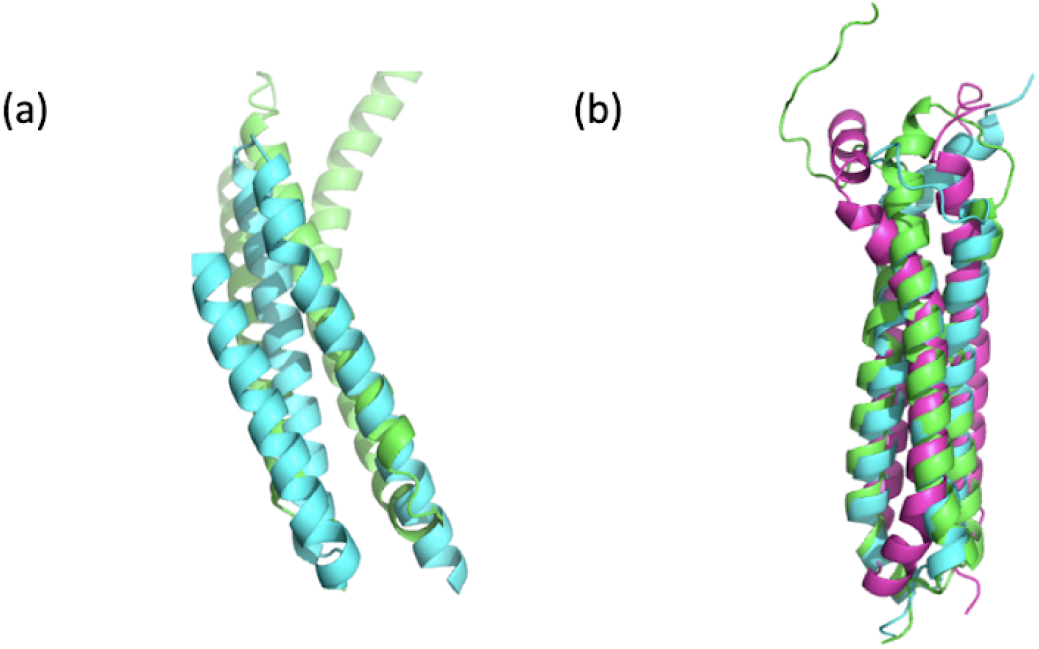
Predicted structure of the hypothetical proteins situated between Coxlll and CtaG in xoxF3 systems from Verrucomicrobia, Gemmatimonadetes and Acidobacteria genomes, (a) Hypothetical protein (green) from RPR_S_B2_Acidobacteria_65_30 was superimposed in pyMOL with the best hit (cyan) PDB:2qyw (RMSD = 2.30) from PDBeFold. (b) Hypothetical proteins from RPR_1_B1_Verrucomicrobia_61_17, RPR_S_A2_Gemmatimonadetes_63_18 and RPR_S_B2_Acidobacteria_65_30 were aligned in pymol (RMSD = 1.170).

**Supplementary Figure 6.**
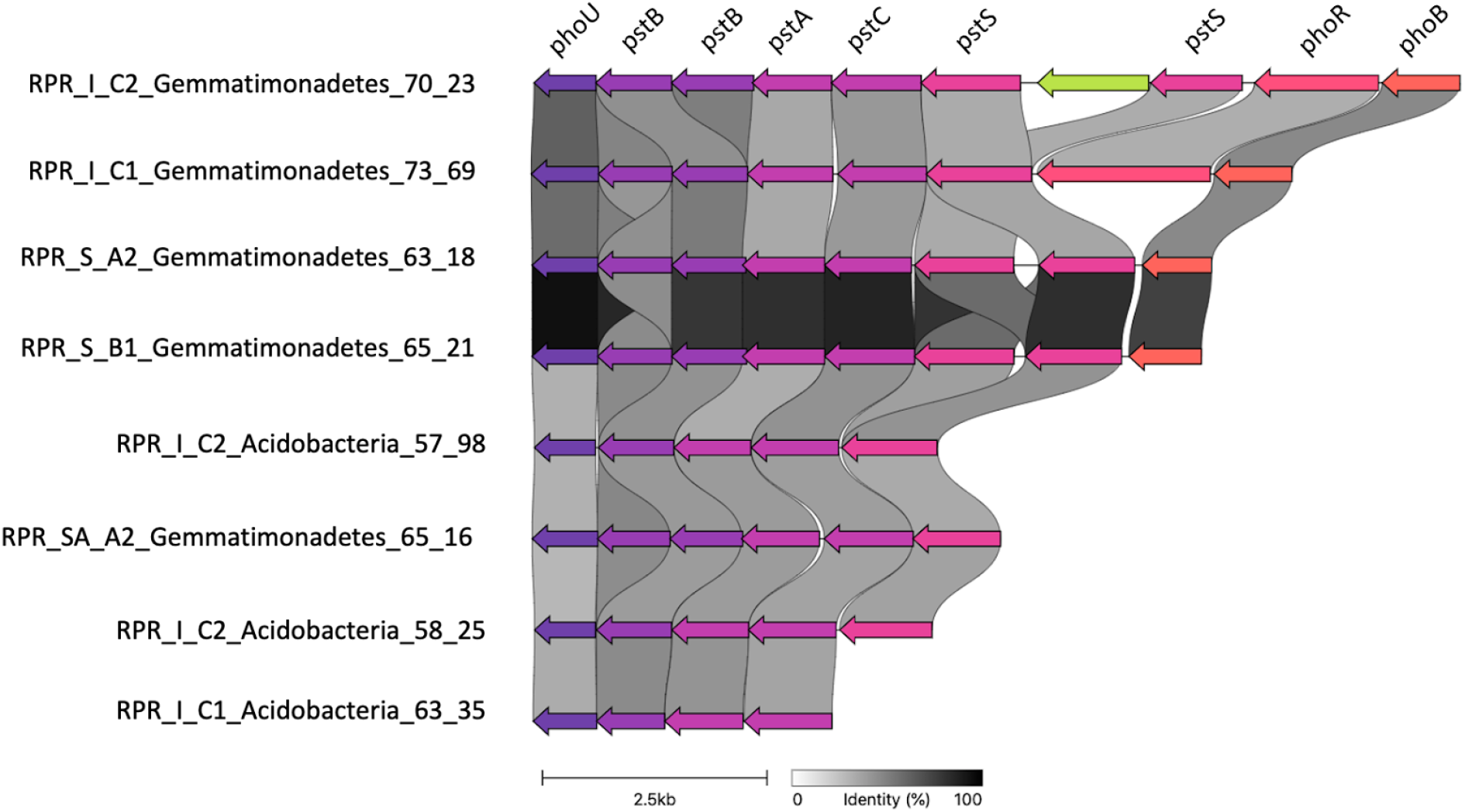
Phosphate regulons partially conserved across Acidobacteria and Gemmatimonadetes genomes recovered from the moderately weathered, highly weathered and soil regions. Grey links indicate the percentage of identity between homologous proteins from different genomes. Abbreviations in order of appearance: pho, phosphate uptake regulon; pst, phosphate specific transporter.

**Supplementary Figure 7.**
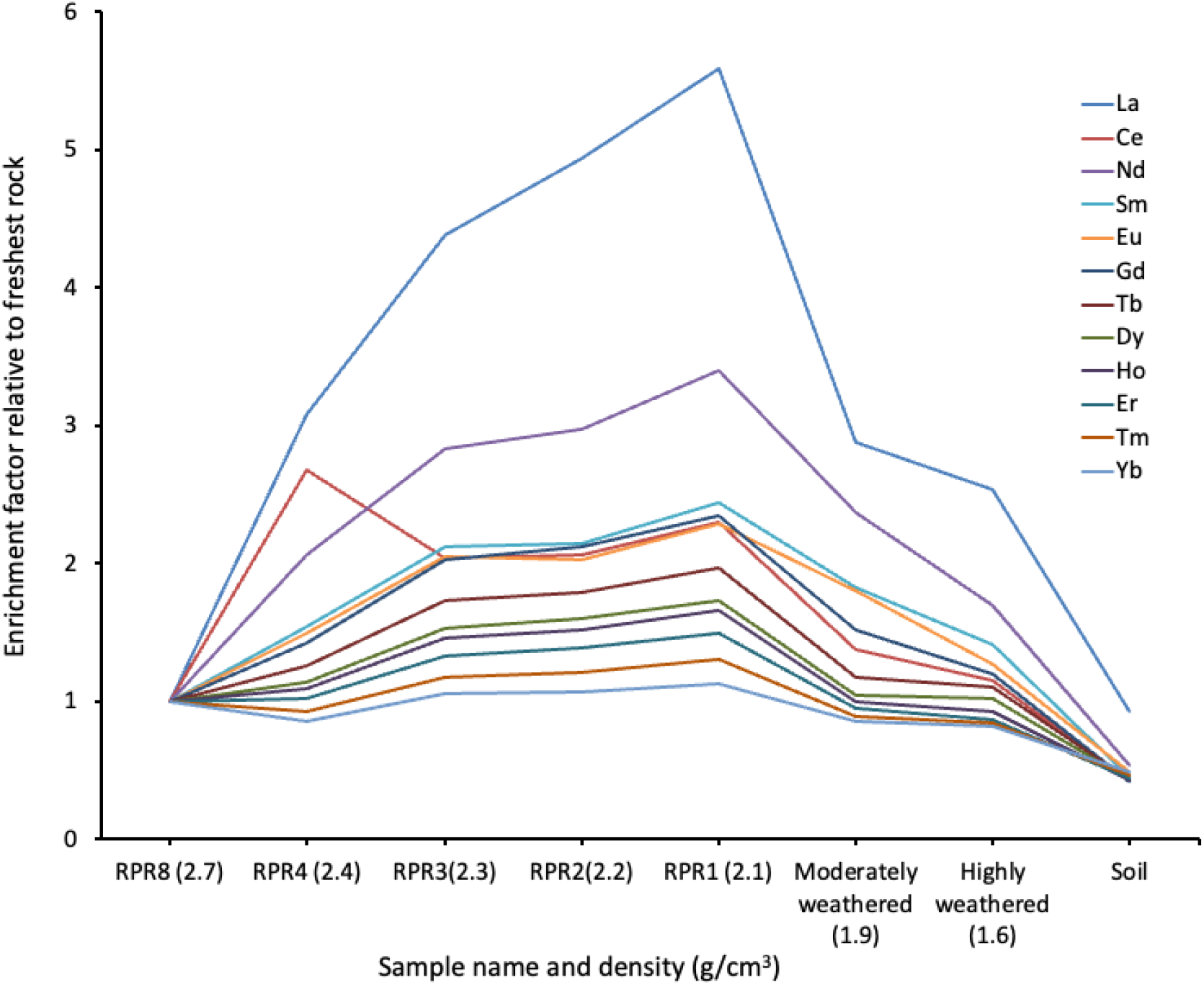
ICP-MS data of all lanthanides and their concentration in the I-type RPR weathered granite profile relative to the freshest material (RPR8). Data are normalised to fresh rock to determine level of REE enrichment and expressed as “enrichment factor”.

**Supplementary Figure 8.**
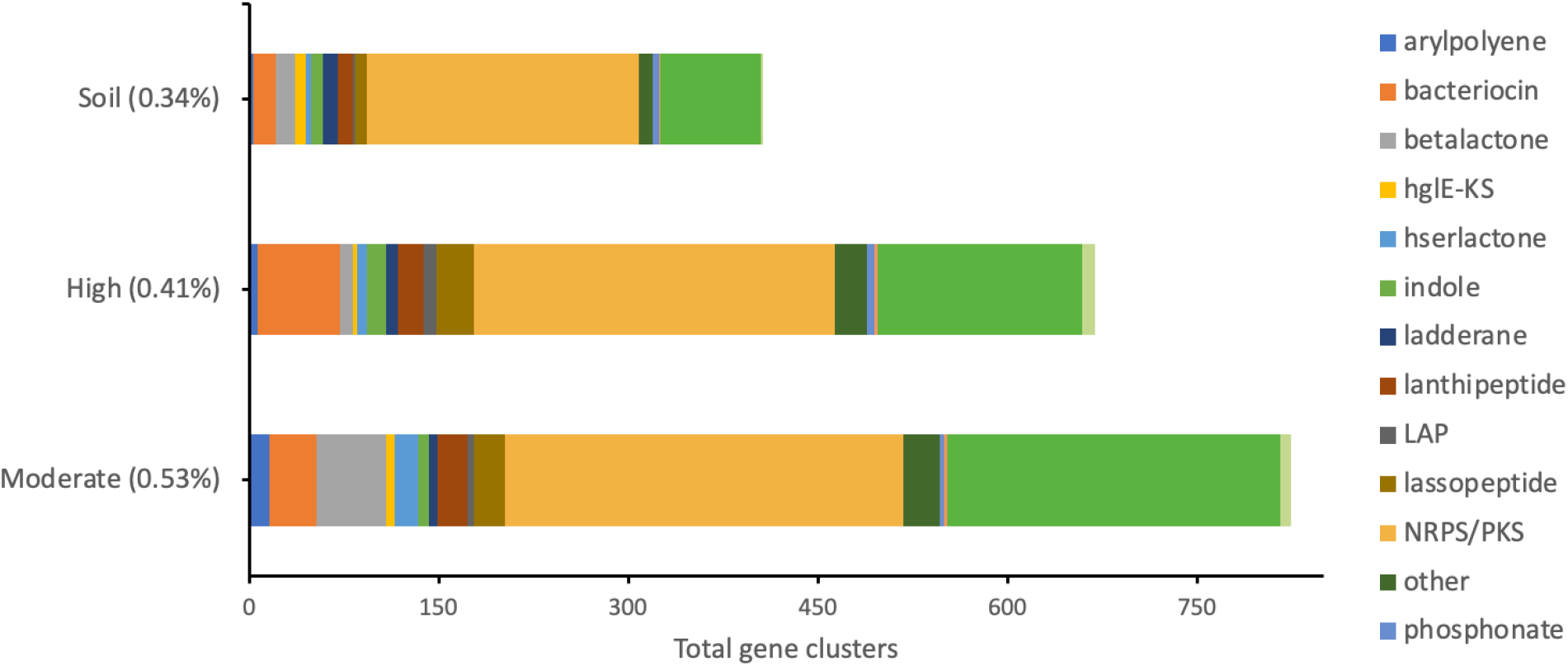
The total number and products of biosynthetic gene clusters (BGCs) predicted in scaffolds greater than 10kb from each metagenome (binned and unbinned sequences) using antiSMASH 5.0. The number of BGCs from each weathered region were combined. BGCs (>10 kb) were normalised to percent abundance in each sample and are displayed beside sample names.

**Supplementary Figure 9.**
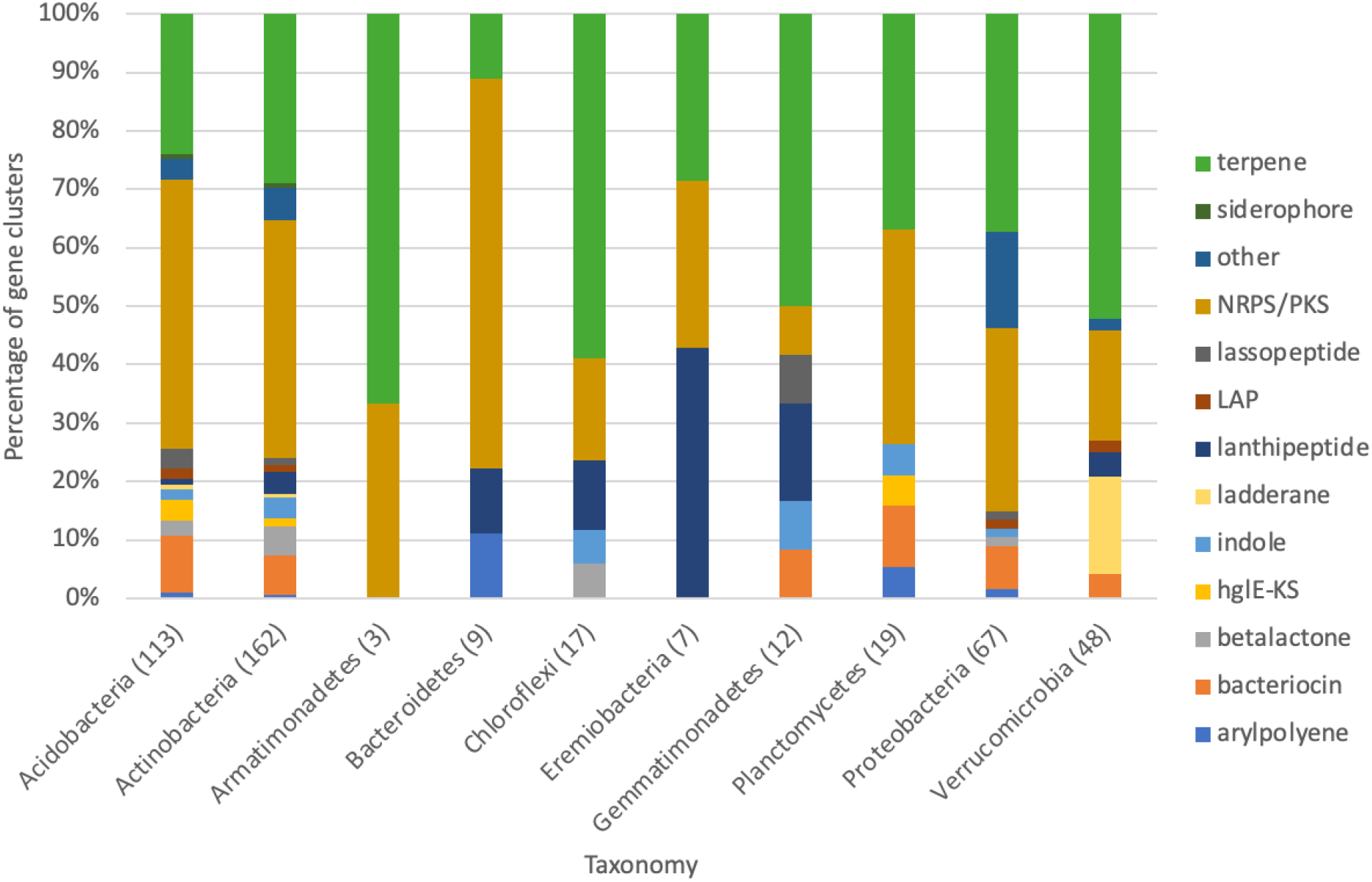
The percentage of biosynthetic gene cluster (BGC) products within taxonomic groups as predicted by AntiSMASH from the dereplicated genomes set. Total number of BGCs in each taxonomy are presented in parentheses beside taxonomies. Only BGCs >10 kb are included.

